# Do morphologically distinct groups correspond to reproductively isolated species? A case study in *Myrmica* ants from Switzerland

**DOI:** 10.1101/2024.12.02.626339

**Authors:** Guillaume Lavanchy, Christophe Galkowski, Kristine Jecha, Anne Freitag, Amaury Avril, Aline Dépraz, Tanja Schwander

## Abstract

The most widely used definition of a species is that it is reproductively isolated from other populations. Yet, most species are described on the basis of morphological criteria, and reproductive isolation is seldom tested. Using the ant genus *Myrmica* Latreille (Hymenoptera, Formicidae) as a model, we ask whether species described as distinct based on (often subtle) morphological differences indeed form reproductively isolated lineages. We collected and morphologically identified 918 *Myrmica* ants from a 3212 km^2^ area in Switzerland. We then combined DNA barcoding (based on COI) and RAD sequencing to identify genetically isolated lineages. Out of the 14 morphological species identified, 13 formed genetically differentiated lineages, while the last one was not supported by our genetic data. Overall, the morphological identification was congruent with genetic lineage delineation for 94.9% of individuals. Our dataset also allowed us to screen for cryptic lineages in the five most frequent species, including in *M. scabrinodis* where cryptic lineages were previously suggested, but we found no evidence for cryptic species. Overall, our results indicate that morphology parallels genetic isolation in the studied species. However, an integrative approach combining morphological identification with nuclear marker genotyping is necessary for confident species identification of all individuals. Finally, our results provide a library of validated COI barcodes for future *Myrmica* specimen identification.

## Introduction

Ever since the publication of Linnaeus’ landmark book, *Systema naturae*, naturalists and biologists have sought to name the organisms they were studying. Yet, if the original idea of classifying all life-forms into discrete categories revolutionized the biological sciences, its application proved to be arduous. The first challenge lies in the definition of the categories themselves as no universal species concept can be uniformly applied across the whole tree of life (Barraclough, 2019; Coyne & Orr, 2004).

The most widely adopted definition of a species among (sexually reproducing) animals is the biological species concept: two populations are said to belong to different species if they do not regularly reproduce under natural conditions (Dobzhansky, 1935). In practice however, assessing reproductive isolation when describing new species is unrealistic. Scientists thus rely on indirect evidence to assess which species their specimens belong to.

The most common indirect line of evidence is morphology (Schlick-Steiner et al., 2010). The underlying assumption is that most of the variation in morphology that can be detected in a set of specimens is species-specific. Yet, this is not necessarily the case. Morphological traits used for species description are not necessarily indicative of reproductive isolation, and individuals in fully inter-breeding populations can differ in morphology because of phenotypic plasticity (Demes et al., 2009; Gruber et al., 2013) or owing to polymorphism at a small set of loci in an otherwise homogeneous gene pool (Cumer et al., 2024; Hager et al., 2022; Lamichhaney et al., 2016; Tuttle et al., 2016; Willink et al., 2024). This may result in over-splitting of lineages that are actually inter-breeding (e.g. Funk et al., 2021). Alternatively, reproductively isolated species may not show detectable morphological variation. Such cryptic species might be frequent in some taxa (e.g. Seifert et al., 2017). In addition, hybrids, which often present intermediate phenotypes, further complicate morphological identification (e.g. Yazdi et al., 2012).

Whether species described on the basis of morphology indeed fit the biological species concept is rarely tested. Yet, given the widespread use of morphological identification for inferring key patterns in ecology and evolution, testing to what extent morphologically distinct groups indeed represent genetically isolated species is crucial. Doing so requires a population genetics approach. Under reproductive isolation, populations can be detected as separate groups that are genetically more similar to each other than to individuals of other species. The integration of morphological criteria is then required to assign genetically distinct groups to taxonomically defined species (integrative taxonomy; Dayrat, 2005).

Here, we ask whether individuals assigned to different species using morphological criteria belong to genetically isolated lineages, focusing on ants of the genus *Myrmica*. The identification of species in many ant genera is challenging, and there is accumulating evidence that hybridization is common in ants (reviewed in Feldhaar et al., 2008). *Myrmica* in particular is a geographically widespread genus for which reliable morphological species identification requires substantial practical experience and comparisons to reference specimens, but genetic studies are scarce and thus far solely based on a few markers (Jansen et al., 2010). We use samples collected during an inventory of the ants of the Canton de Vaud, in Switzerland (Szewczyk et al., 2024). This 3212 km^2^ area ranges from the Jura mountains to the Alps, separated by a lowland region, and encompasses diverse habitats. Samples were collected by means of a citizen science project combined with structured sampling. We identify *Myrmica* samples to the species level using classical morphological criteria (Seifert, 2018) and use a combination of genome-wide SNP genotyping (via RAD sequencing) and COI barcoding to define genetically distinct clusters of individuals. We then compare the species detected by using solely morphological identification versus solely genetic criteria. We also take advantage of our dataset consisting of 692 genotyped *Myrmica* to screen for cryptic species. In particular, we evaluate the presence of a cryptic species in *M. scabrinodis* as suggested by Ebsen et al. (2019), based on the occurrence of two mitochondrial lineages. In addition to assessing the extent to which morphology-based descriptions fit the biological species concept, we provide a library of curated barcodes for thirteen central European species of *Myrmica*, which will be an invaluable resource for future species inventories and monitoring.

## Material and Methods

### Study area and sampling

We studied all *Myrmica* samples collected during a sampling campaign in the *Canton de Vaud*, Switzerland in 2019. The sampling campaign combined two sources: a citizen science project during which members of the public collected samples from anywhere in the focal region (described in Avril et al., 2019) and a structured sampling (Szewczyk et al., 2024). The latter consisted of 44 sites (1 km^2^ each) placed on a regular grid across the study area. At each site, we searched for ant nests (colonies) at 25 plots selected in a random stratified manner, proportional to the frequencies of different habitats at each site. For both the citizen science and structured sampling approaches, about 10 workers of each detected nest were collected. Samples were stored in 70% EtOH until DNA extraction. The location of all 918 *Myrmica* samples obtained from the two sampling approaches is shown in Figure 1.

**Figure 1:**
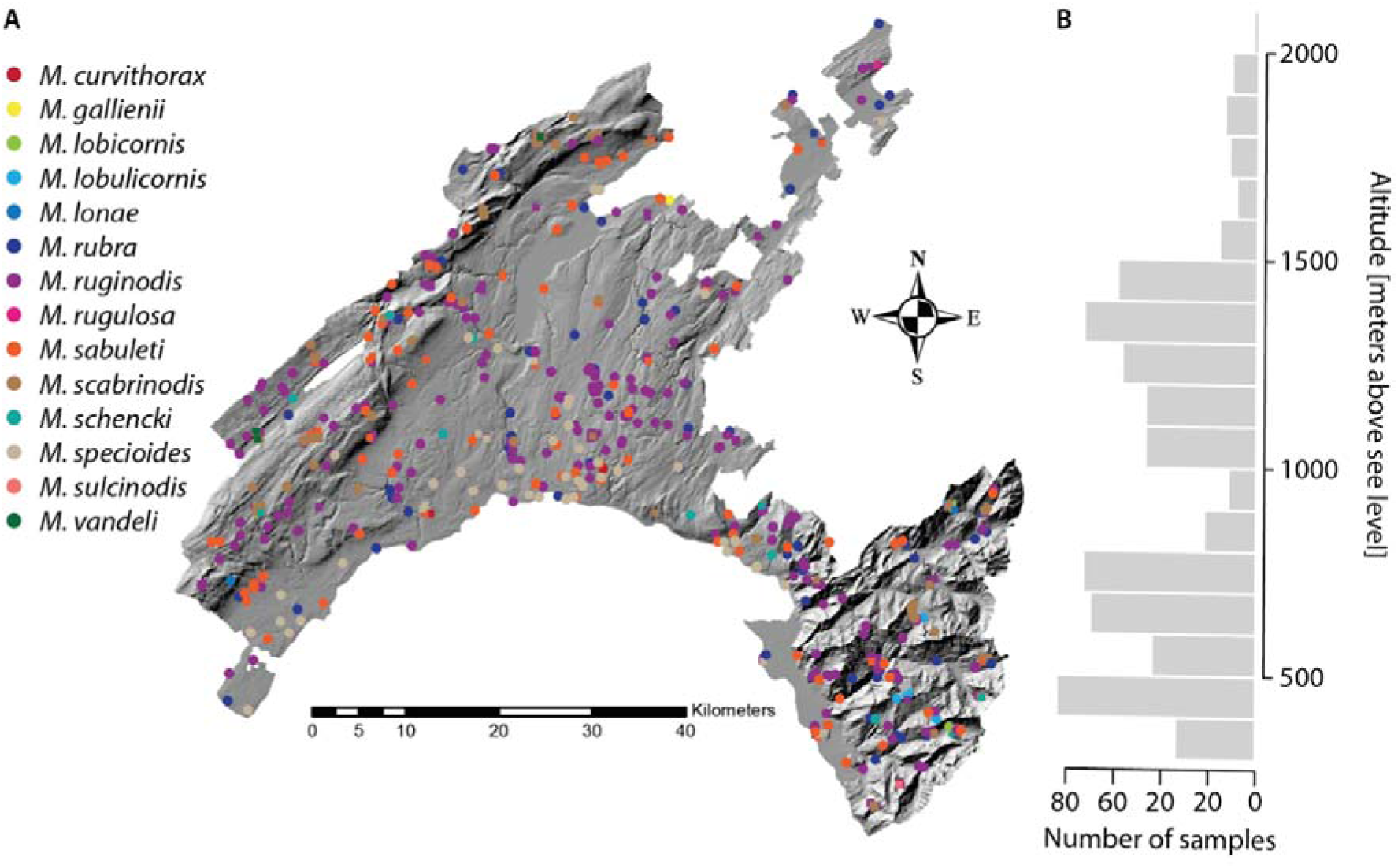
**A**: Map of the study area showing the location of all sampled *Myrmica* nests. The Jura mountains are located in the western part of the canton, separated from the Alps in the southeast by the Plateau lowlands. Points are coloured according to *a priori* species identification based on morphology following Seifert (2018).

Each of the 918 *Myrmica* samples was identified morphologically to the species level based on scape and petiole shape and propodeal spine length, following Seifert (2018). We then selected samples for genetic analyses. For all except one morpho-species we used all available samples. For the most frequent morpho-species, *M. ruginodis,* we used all 134 samples from the structured sampling and only 126 out of 260 citizen science-based samples to avoid geographically redundant samples. The total number of samples analyzed across all species was 765. All voucher specimens are deposited at the Museum of Zoology, Lausanne, Switzerland.

### Wet lab and sequencing

We haphazardly selected one worker per sample for genetic analyses. DNA was extracted from three legs. DNA extraction and COI barcoding was performed by the Canadian Center for DNA Barcoding. A fragment of the COI mitochondrial gene was amplified using primers LepF1 and LepR1 (Hebert et al., 2004). The specific protocols used for extraction, PCR amplification and sequencing are available at https://ccdb.ca/resources (last checked 14.07.2022).

In addition, we generated RAD sequencing data for all individuals using the protocol described in Brelsford et al. (2016). Briefly, we digested genomic DNA with restriction enzymes EcoRI and MseI, ligated custom barcoded adapters, amplified the resulting fragments in 16 PCR cycles and selected fragments ranging between 280 and 430 bp using 2% agarose cassettes on a BluePippin (Sage Science). The resulting libraries were sequenced on a total of eight Illumina HiSeq 2500 lanes (single end, 150 bases).

### SNP calling

We built SNPs using *stacks* v2.3e (Catchen et al., 2011). We first demultiplexed the reads using the *process_radtags* command with the options *-c -q -r -t 143*. In order to prevent putative contaminations from influencing our conclusions, we discarded reads corresponding to other ant genera using the competitive mapping approach developed by Jecha et al. (2024). Briefly, we mapped the reads to a concatenated file containing other local ant genera that were studied in the same laboratory at the same period. We retained only reads that mapped preferentially to the genome of *Myrmica rubra* from Romiguier et al. (2022). We then mapped the filtered reads onto the genome assembly of *Myrmica rubra* using the *mem* algorithm of *bwa* version 0.7.17 (Li & Durbin, 2009). We used *samtools* version 1.4 (Li et al., 2009) to convert mapping files into bam format and to generate summary statistics. We then built loci using *gstacks*, enabling the options *--phasing-dont-prune-hets* and *--ignore-pe-reads*. Finally, we ran *populations* with the option *--write-single-snp* to retain only one SNP per RAD locus.

We filtered the data in vcftools 0.1.14 (Danecek et al., 2011). We retained genotype calls with a minimum depth of 8, and loci for a maximum locus-wide average depth of 200, a minor allele count of two and presence in at least 75% of individuals. We discarded all individuals which had over 75% of missing data (n=51).

We further filtered our dataset for potential contaminations following the strategy developed by Jecha et al. (2024). This strategy is based on allelic depth ratio (ADR), i.e. the ratio of reads supporting the two alleles in heterozygotes. In diploid organisms (such as worker ants), both alleles are expected to be supported approximately by the same number of reads in real heterozygotes. However, if a heterozygous genotype is caused by a contamination, since the amount of contaminated DNA is often lower than the real sample, ADR is expected to be biased towards the real allele. We estimated the expected distribution of ADR based on observed coverage at heterozygous alleles (see Jecha et al., 2024 for details). We then discarded 44 individuals whose median ADR was beyond the 95% quantile of the expected distribution. We also corrected the genotypes of retained individuals when the ADR of individual genotypes was beyond the 75% quantile of the distribution, by discarding the allele with the lowest number of reads. Our final dataset contained 6924 SNPs and 666 individuals. We generated datasets for subsets of individuals for subsequent analyses by filtering with the same parameters.

### COI phylogeny

We reconstructed the COI phylogeny of all our samples, including published sequences from other studies to cover genetic diversity within species better. We included all *Myrmica* sequences from Blatrix et al. (2020), Jansen et al. (2009) and Leppänen et al. (2013). We could not include individuals from Ebsen et al. (2019) as the region of COI they amplified did not overlap with ours. We did not include any additional sequences from Genbank because of uncertain morphological identifications. We aligned the sequences using the *L-INS-i* algorithm of *mafft* v7.475 (Katoh et al., 2005), with the mitochondrial genome of *M. scabrinodis* (GenBank accession number NC_026133; Babbucci et al., 2014) as reference for alignment. We then built a Maximum Likelihood tree using IQ-TREE v2.0.6 (Nguyen et al., 2015) using ModelFinder Plus (Kalyaanamoorthy et al., 2017) to select the best substitution model, and assessed its robustness with 1000 UltraFast Bootstraps.

### Genetic species delineation

In order to delineate species solely based on genetic information, we identified genetically isolated groups (“clusters”). We used two complementary approaches to identify genetic clusters. First, we ran *admixture* (Alexander et al., 2009) with k ranging from 1 to 20, in 10 replicates for each value of K. Based on the cross-validation error, the best value of K was seven (Figure S1). We then selected the run with the lowest CV error. One drawback of admixture and related approaches is that the results are highly dependent on sampling (Ravagni et al., 2021; Shringarpure & Xing, 2014). These approaches will for example tend to oversplit groups with large sample sizes, but fail to identify groups with smaller sample sizes and consider them as admixed between other groups. For this reason, we used a second approach based on multivariate analyses. Specifically, we performed distance-based MultiDimensional Scaling (MDS) on the SNP dataset containing all individuals using the *isoMDS* function implemented in the R package *MASS* (Venables & Ripley, 2002). MDS is a way to visualize pairwise distances between individuals that is robust to missing data. Because distances between closely related species tend to be masked by a few distant species, we performed several MDS with different subsets of the data. We first included all individuals to define major groups, and then looked for structure within each group by performing a new MDS only with individuals from that group. Finally, we screened for potential cryptic lineages within genetically well-defined species by performing an MDS for each species with at least 50 samples.

## Results

Morphological identification to the species level was possible for 661 out of the 666 successfully genotyped samples and suggested the presence of 13 species. The remaining 5 samples (0.7%) could not be confidently attributed to a species based on morphology. The 13 morpho-species had highly variable abundances, with 92.6% of the samples belonging to only five different species and five species represented by fewer than four individuals (Table 1). A 14th species, *M. lonae*, was detected morphologically, but was not validated genetically (see below).

**Table 1:** Comparison of the results of the morphological identification (rows) and final identification based on the combination of all available information (“integrative identification”). Bold numbers correspond to matching morphological and integrative identification.

|  |  | Integrative identification |  |  |  |  |  |  |  |  |  |  |  |  |
| --- | --- | --- | --- | --- | --- | --- | --- | --- | --- | --- | --- | --- | --- | --- |
|  |  | *M. curvithorax* | *M. gallienii* | *M. lobicornis* | *M. lobulicornis* | *M. rubra* | *M. ruginodis* | *M. rugulosa* | *M. sabuleti* | *M. scabrinodis* | *M. schencki* | *M. specioides* | *M. sulcinodis* | *M. vandeli* |
| Morphological identification | *M. curvithorax* | **2** |  |  |  |  |  |  |  |  |  |  |  |  |
|  | *M. gallienii* |  | **1** |  |  |  |  |  |  |  |  |  |  |  |
|  | *M. lobicornis* |  |  | **4** | **1** |  |  |  |  |  |  |  |  |  |
|  | *M. lobulicornis* |  |  | **3** | **15** |  |  |  |  |  |  |  |  |  |
|  | *M. rubra* |  |  |  |  | **83** | **1** |  | **2** |  |  |  |  |  |
|  | *M. ruginodis* |  |  |  |  | **5** | **228** |  |  |  |  |  |  |  |
|  | *M. rugulosa* |  |  |  |  |  |  | **1** |  |  |  |  |  |  |
|  | *M. sabuleti* |  |  |  |  |  |  |  | **129** | **7** |  |  |  |  |
|  | *M. scabrinodis* |  |  |  |  |  |  |  | **3** | **104** |  | **1** | **1** |  |
|  | *M. schencki* |  |  |  | **3** |  |  |  |  |  | **12** |  |  |  |
|  | *M. specioides* |  |  |  |  |  |  |  |  | **1** |  | **48** |  |  |
|  | *M. sulcinodis* |  |  |  |  |  |  |  |  |  |  |  | **2** |  |
|  | *M. vandeli* |  |  |  |  |  |  |  |  |  |  |  |  | **3** |
|  | *M. lonae* |  |  |  |  |  |  |  | **1** |  |  |  |  |  |
|  | unknown |  |  |  | **1** |  | **1** |  | **3** |  |  |  |  |  |

The genetic species delineation revealed the presence of seven distinct nuclear genetic clusters (Figure 2C). The abundances of the different clusters were also variable (range: 3 - 233 individuals per cluster). Contrary to the morphological identification, the genetic species delineation alone did not detect very rare species, as 17 individuals from rare species were inferred to be admixed between the more frequent lineages.

**Figure 2:**
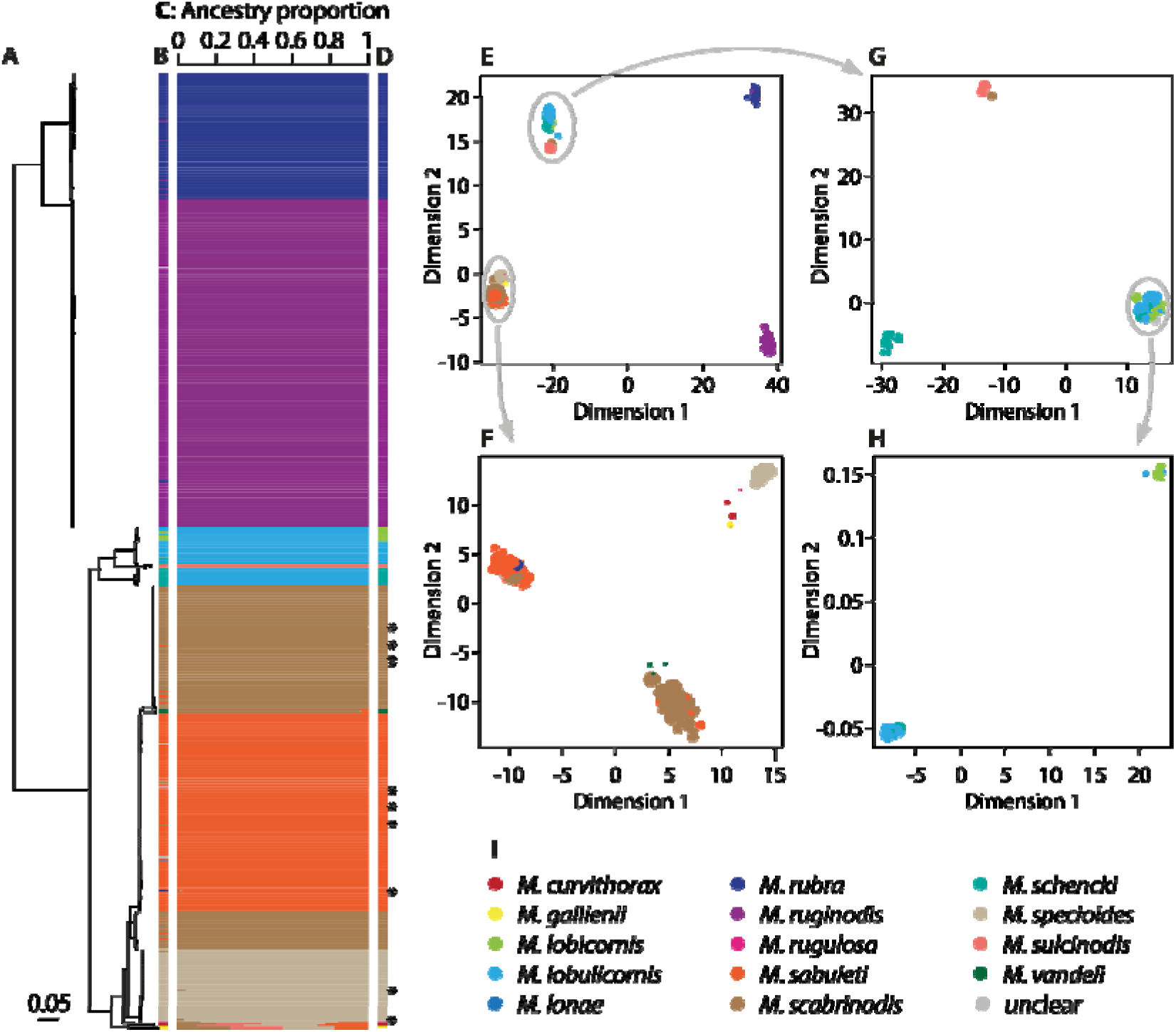
Correspondence between morphological and genetic species delineations. **A**: mtDNA phylogeny based on 613 bases of COI. **B**: morphological identification. **C**: Ancestry estimation from admixture. The barplot shows the inferred contribution of each of k = 7 clusters to each individual. Note the apparent hybrid ancestry inferred by admixture for the species *M. curvithorax, M. gallieni* and *M. sulcinodis* (bottom) which is caused by the rarity of these species in the dataset rather than by true admixture (see text for details). **D**: Final species assignment based on the combined information. Individuals denoted with stars were inferred to be true hybrids. **E**: MDS plot of all individuals. Dots are coloured based on their morphological identification. **F-H**: MDS plots of subsets of individuals. Dot size in **E - H** proportional to heterozygosity. Note that in **G** and **H**, artificial noise was added to the data to separate the points to improve display. **I**: Colour code.

We then assessed the correspondence between morphological and genetic species delineation. We used the genetic groups identified using the combination of nuclear and mitochondrial data (Figure 2) and assigned a species name using the morphological identification and published COI sequences (Figure S2). There was a very high congruence between morphological and genetic species delineation for five out of the 14 *Myrmica* species (*M. rubra*, *M. sabuleti, M. ruginodis*, *M. scabrinodis* and *M. specioides*). Each of these species formed a single nuclear cluster and the morphological identification matched in 95.9% of individuals belonging to these clusters (Table 1). Four of these five species also formed monophyletic mtDNA clades. The fifth (*M. scabrinodis*) consisted of two polyphyletic mtDNA clades, which likely indicates incomplete lineage sorting for mitochondria as has been observed in other ants (see also below). The success of the morphological identification was more variable for the rare species, with 100% accuracy in some species (*M. curvithorax, M. gallienii, M. schencki, M. vandeli*), and a low proportion of accurate determinations in others (*M. lobicornis*: 4/7; *M. sulcinodis*: 2/3). The detailed comparisons between morphological and genetic species delineations for each ambiguous species are described separately in the Supplementary material.

The morphological identifications were congruent with the identification based on the combination of all evidence (i.e. integrative approach) for 632 (94.9%) individuals, while the two approaches led to different results for 34 (5.1%) individuals. Correspondence between the original morphological identification and the final species assignment based on the genetic information are summarized in Table 1.

As described above, the integrative approach also allowed us to identify 17 individuals which featured a signature of admixture (Figure 2C). Seven had rare mitochondrial DNA and were identified morphologically as rare species, consistent with rare species being identified by admixture as a mix between frequent lineages. The remaining 10 individuals with signs of admixture had the same mitochondrial lineage as the cluster to which they were most similar, corroborating that they were hybrids. None of them looked like F1 hybrids, and all had ancestry proportions of their main genetic cluster > 93.75% (which is expected from hybrids having undergone at least three generations of backcrossing).

Finally, we used our dataset to investigate the presence of cryptic species. Contrary to the suggestion of Ebsen et al. (2019), there was no evidence for cryptic species in *M. scabrinodis*. All individuals morphologically and genetically assigned to *M. scabrinodis* belonged to two polyphyletic mtDNA lineages, probably corresponding to the two haplotypes found by Ebsen et al. (2019). Their position on the phylogeny relative to *M. sabuleti* suggests that the frequent haplotype (top on Figure 2) corresponds to lineage A in Ebsen et al. (2019), while the rare haplotype (bottom on Figure 2) corresponds to lineage B (Figure 3A, B). It was impossible to verify this formally as they sequenced a fragment of COI that does not overlap with the commonly-used barcode region. The frequent haplotype was found throughout our study area, while the rare one was found only in the Jura mountains, in the western part of our study area (Figure 3C). Individuals of the two mitochondrial lineages formed a single cluster in admixture (Figure 2C). However, the two clades were slightly differentiated and on the MDS (although with significant overlap; Figure 3D).

**Figure 3:**
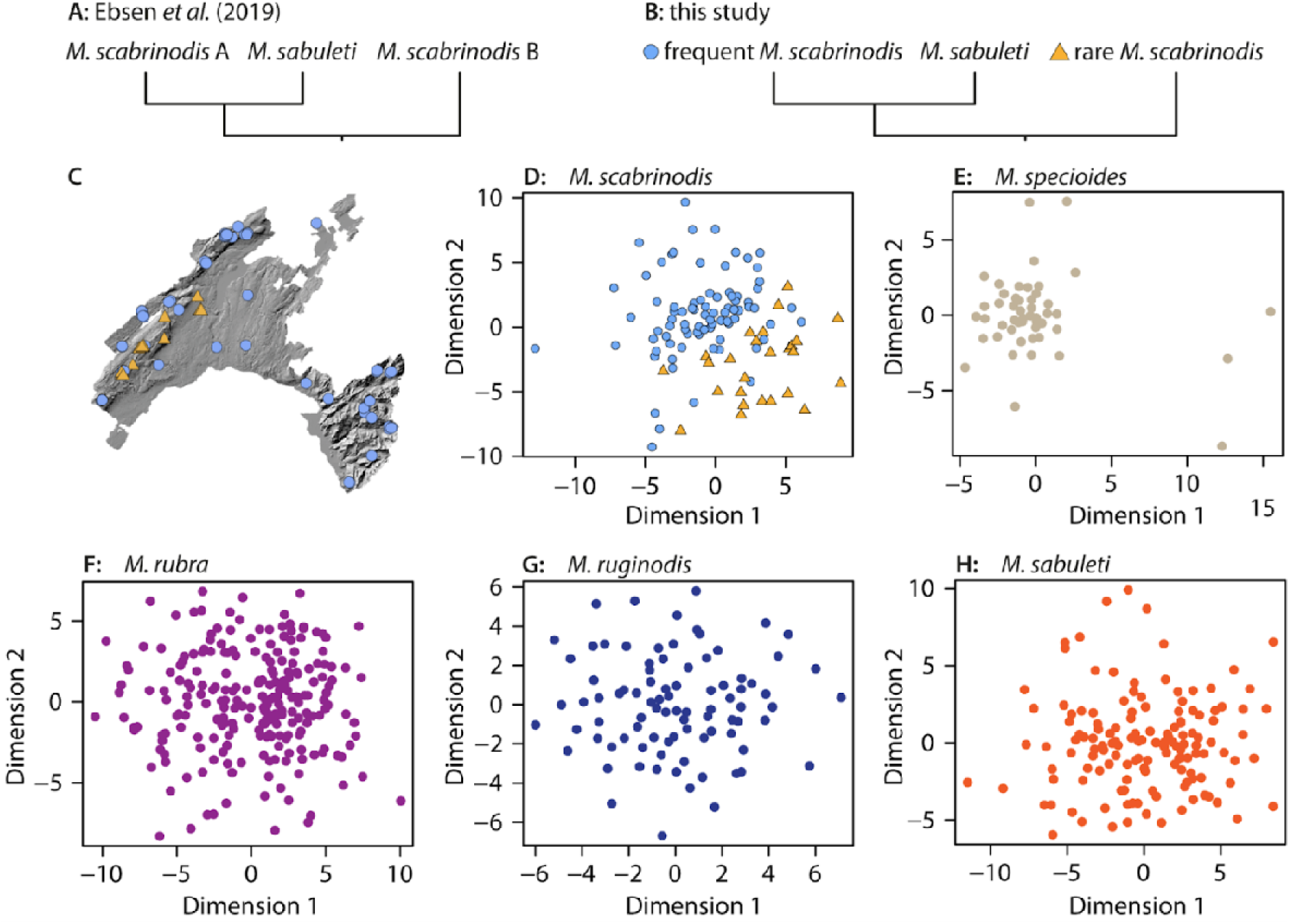
Assessing the presence of cryptic species. **A**: Cladogram of the relationships between the two *M. scabrinodis* haplotypes of Ebsen et al. (2019) and *M. sabuleti*. **B**: Cladogram of the relationships between the two *M. scabrinodis* mitochondrial haplotypes found in this study and *M. sabuleti* (simplified from Figure 2A) **C**: Map of study area with location of individuals belonging to the two mitochondrial haplotypes. **D**: MDS plot of all *M. scabrinodis*. Colours as in **B**. **E**: MDS plot of all *M. specioides*. **F**: MDS plot of all *M. rubra*. **G**: MDS plot of all *M. ruginodis*. **H**: MDS plot of all *M. sabuleti*.

In addition to *M. scabrinodis*, we tested for the presence of cryptic lineages in the four remaining species with at least 50 samples (*M. ruginodis, M. rubra, M. sabuleti, M. specioides*). In these three species, all samples formed one homogeneous group, which indicates that there are no cryptic lineages in our study area (Figure 3E – H).

## Discussion

Morphological identification is traditionally used to discriminate species for monitoring and inventories. It is still often used as the main line of evidence when describing new species. However, the extent to which morphological differentiation parallels genetic isolation is unknown in many taxa. As a result, morphologically distinct groups can bear little evolutionary and ecological relevance. Relying on them to measure biodiversity might overestimate diversity by mistaking plastic morphological variation with species differences, or underestimate it by overlooking cryptic species. In addition, hybridization can result in introgression of morphological characters, further blurring species boundaries. Testing whether morphological differences reliably discriminate genetically isolated groups is thus crucial.

Here, we tested whether morphologically distinct groups indeed represented true species in 692 *Myrmica* ants found in the *canton de Vaud*, Switzerland. Species from this genus were described based on morphological criteria and genetic confirmation is scarce: the phylogeny of Jansen *et al*. (2010) based on a few genes in one individual per morphological species suggested that they differed genetically, but all other studies (e.g., Blatrix et al., 2020; Ebsen et al., 2019; Leppänen et al., 2013) were conducted only on mitochondrial DNA. Overall, we found strong evidence that the six most common morphological species in our samples each correspond to a homogeneous genetic group that is isolated from others, indicating that they are true species. Another seven morphological species were also likely reproductively isolated species, but larger sample sizes would be needed to draw definitive conclusions.

Our analyses did not detect any cryptic species, even where previously suggested: genetic differentiation at the nuclear level between the two mitochondrial haplotypes in *M. scabrinodis* that Ebsen *et al*. (2019) interpreted as a putative cryptic species was in fact very low, with much overlap. This low level of differentiation is unlikely to deserve the subspecies label and more likely reflects postglacial recolonization from different refugia or incomplete lineage sorting. Sampling throughout the range of the species would be necessary to confirm this hypothesis. Such polymorphism has already been observed in other ants (e.g. *Tetramorium*; Wagner et al., 2017). The frequent *M. scabrinodis* mitochondrial haplotype was found throughout our study area. The rare one was found only in a restricted area along the Jura mountain range, where it was sympatric with the frequent one. Ebsen et al. (2019) also found that their two haplotypes co-occurred across parts of their range, although both were found to be widespread throughout Europe. The mosaic distribution of the two haplotypes could then be due to local extinction of one haplotype.

We found no evidence that our sample identified as *M. lonae* represents a distinct species. The only individual identified morphologically as *M. lonae* belonged to *M. sabuleti*. This result, together with the apparent polyphyly of *M. lonae* mitochondrial haplotypes within *M. sabuleti* (Ebsen et al., 2019), raises the question of whether *M. lonae* could be a morphologically distinct form of *M. sabuleti*, with no genetic isolation from that species. More extensive sampling would be needed to evaluate the different hypotheses.

Our findings that morphological identification matched the integrative identification in 94.9% of cases shows that morphology is generally representative of true species for the studied *Myrmica*. Morphological identification of rare species (represented by only a few samples) was often less reliable. This could be due to rare species being underrepresented in collections, which would result in a poor coverage of intraspecific phenotypic variation.

Our results indicate that morphology-based specimen identification can be used reliably for monitoring of the most frequent species of *Myrmica* in our study area. We expect that the low error rate for these species should not influence quantitative conclusions in a significant way. However, morphological identification might not be sufficient for some of the rare species, or when the identity of each specimen is of interest. Complementing identification based on qualitative trait estimation with quantitative morphological methods more robust to subjective judgements such as Numeric Morphology-Based Alpha-Taxonomy (NUMOBAT, Seifert, 2009) could address some of these issues. However, NUMOBAT is time-consuming and still relies on the assumption that the measured phenotypic traits are fixed between species.

One alternative to morphological identification is DNA barcoding. Sequencing always the same locus (in the case of animals, a fragment of the mitochondrial gene COI) provides a large database of reference individuals with which to compare new specimens. Barcoding has become very cost-effective, can be performed with as little material as a single ant leg, and can be performed in a very high throughput manner using amplicon sequencing. However, it relies on the assumptions that all species have different COI haplotypes, and that all haplotypes found in one species form a monophyletic group. Yet, these assumptions are rarely met. Funk & Omland (2003) found that about one quarter of arthropods have polyphyletic mitochondrial DNA (mtDNA), although part of their results can also be explained by cryptic species. In ants in particular, cryptic species (Seifert, 2009), mtDNA polyphyly (e.g., Wagner et al., 2017) and lack of mtDNA polymorphism between species (Jansen et al., 2009) are commonplace. For these reasons, barcoding can only enable reliable specimen identification once a valid barcode library has been constructed for a given geographic region by validating the barcodes via integrative taxonomy. Here, we showed that barcoding would correctly identify the 13 species of *Myrmica* in the surveyed region. Our barcodes can be used as a reference to identify future specimens collected in the same area. Specimens collected outside this region with barcodes similar to ours are likely the same species, but one cannot exclude divergent lineages without COI polymorphism (for example owing to introgression).

We found 10 individuals (1.5%) showing signs of admixture. This is higher than the average estimated proportion in most animals (Mallet, 2005). This can be due to the power of our genomic data to detect low levels of admixture (after several generations of backcrossing). Alternatively, it may be in line with previous suggestions that hybridization might be particularly frequent in ants (Feldhaar et al., 2008; Seifert, 1999; Umphrey, 2006; Weyna et al., 2022). The causes of this frequent hybridization are unknown. Mating swarms of several *Myrmica* species are known to overlap both in phenology and location (Woyciechowski, 1987, 1990b). In addition, (Woyciechowski, 1990a) observed that gynes of *M. rubra* were not choosy among conspecific males. Whether this also applies to males of other species is not clear, but studies on other ants have shown that gynes preferentially mate with conspecific males (Blacher et al., 2022).

Overall, our results indicate that morphology does reflect reproductive isolation to a large extent in the studied species of *Myrmica*. Combining genomics with morphology and barcoding enabled us to validate described species, screen for cryptic species, and reject a previously hypothesized cryptic species in *M. scabrinodis*. Our validated barcodes can be used as references for future specimen identification. Replicating our approach to other taxonomic groups will allow to verify to what extent morphology is a universal proxy for reproductive isolation, and in which taxa it can be used reliably for species description.

## Acknowledgements

We are grateful to the 608 people who collected ants as part of the “Opération fourmis” citizen-science inventory, and Marc Bastardot, Marjorie Labédan and Simon Vogel for their help with processing the samples.

This study was supported by funding from a bequest of M. Rullens to the University of Lausanne, the fondation Herbette, the Société Vaudoise des Sciences Naturelles and the Retraites Populaires.

## Author contribution

GL: Conceptualization, Formal analysis, Investigation, Visualization, Writing – original draft; CG: Investigation; KJ: Formal analysis; AF: Data curation, Writing – review & editing; AA: Data curation; AD: Data curation; TS: Conceptualization, Data curation, Funding acquisition, Supervision, Writing – review & editing.

## Data and code availability

All code used for the analyses is available on GitHub: https://github.com/glavanc1/Myrmica_morpho_geno/.

All demultiplexed reads are archived on SRA (Bioproject PRJNA1188312).

COI sequences for all confidently identified samples are available on Genbank (accessions PQ620386 - PQ621039; accession number of each sample is available in Table S1) and the alignment with all samples used for the phylogeny is available on GitHub.

