## Supplementary figures and images for "Do morphologically distinct groups correspond to reproductively isolated species? A case study in *Myrmica* ants from Switzerland"

### Figure S2

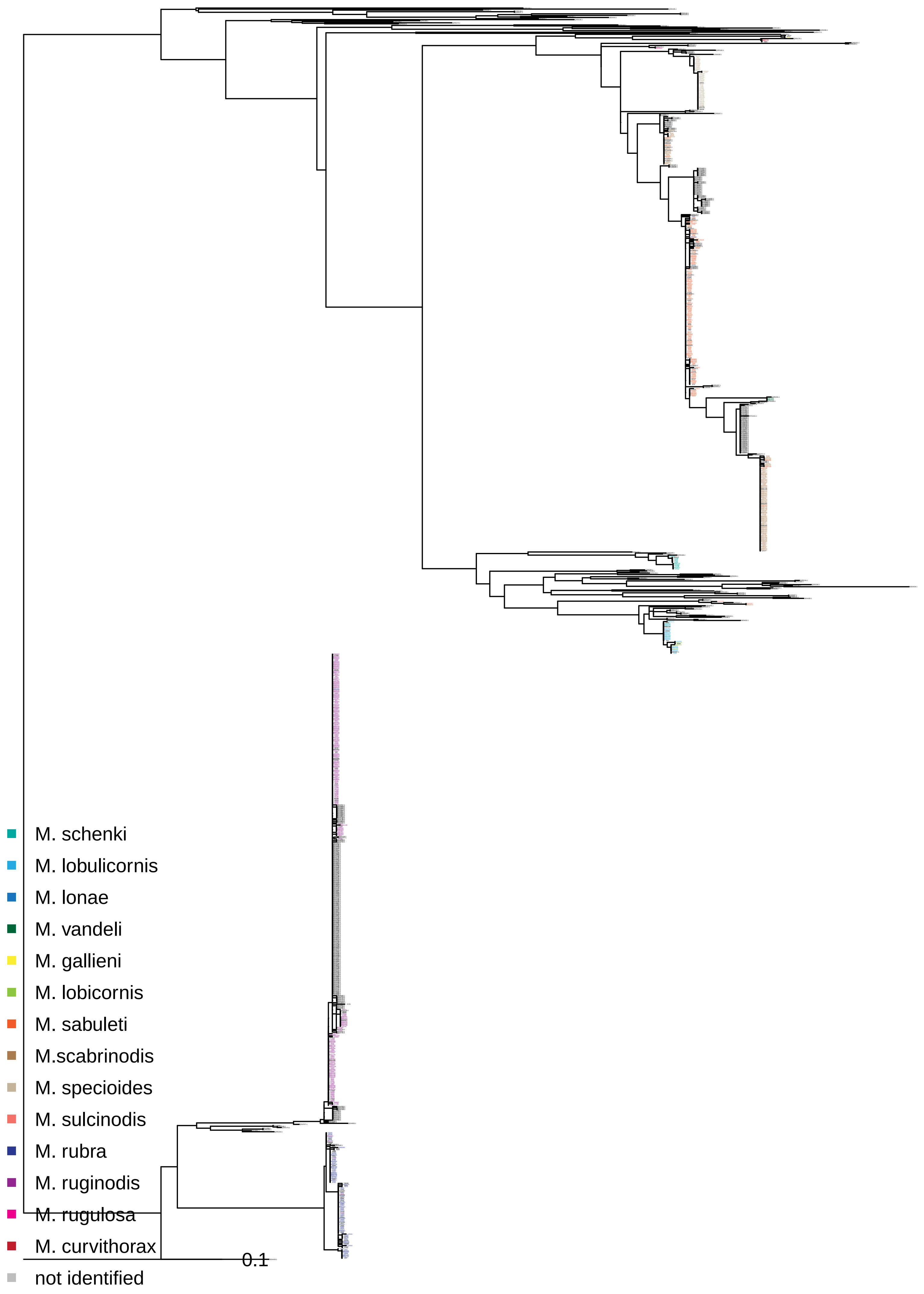
