## Supplementary Material for "Do morphologically distinct groups correspond to reproductively isolated species? A case study in *Myrmica* ants from Switzerland"

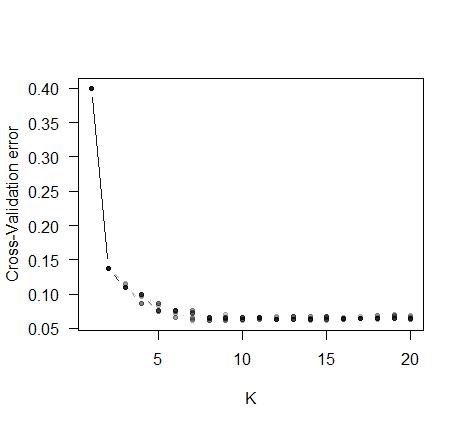

**Figure S1**: Cross-Validation error for the different admixture runs. A plateau is reached around K = 7.

**Figure S2 legend**: mtDNA tree with all published sequences. Individuals names are coloured according to morphological species identification. Individuals in black are reference individuals, and their names correspond to their GenBank accession numbers.

Supplementary text

For each species or group of closely related species we compared the assignment of individuals based on morphological and genetic criteria. We here detail these comparisons for the ambiguous species or specie pair; the summaries for all species are included in the main text and Table 1.

*M. lobicornis* and *M. lobulicornis*

Twenty-one out of 23 individuals identified morphologically as *M. lobicornis* or *M. lobulicornis* formed one cluster in admixture and on the MDS. They formed one monophyletic mtDNA clade containing two sub-clades with very shallow divergence (0.2%; comparable to within-species diversity in other species). The first sub-clade contained a published sequence of *M. lobicornis* and four out of the five individuals identified morphologically as *M. lobicornis*, together with three individuals morphologically identified as *M. lobulicornis*. The second sub-clade consisted of 15 out of 18 individuals identified as *M. lobulicornis*, together with three *M. schencki*, one *M. lobicornis* and one individual for which morphological identification was inconclusive. However, it contained no published sequence, and the only available published sequence of M. lobulicornis was related to a different clade (Figure S2). Our MDS plot on individuals of these two clades revealed that they indeed formed two separate nuclear clusters (Figure 2H). We therefore conclude that the first genetic cluster most likely corresponds to *M. lobicornis* and the second to *M. lobulicornis*, and that the two species are genetically very similar.

The five remaining species identified based on morphology were represented by very few samples (between one and three per species). Four of them likely belong to genetically divergent groups, although this is difficult to corroborate with the limited samples available.

*M. curvithorax*

Three samples were identified morphologically as *M. curvithorax*, but we obtained nuclear data for only two of them. These two clustered in the MDS and showed the same signature of mixed ancestry in the admixture results. There was no published sequence of *M. curvithorax*, but the closest published sequence was from *M. salina*, which has been mistakenly used as a synonym for *M. curvithorax* (Seifert 2018). We conclude that they are indeed *M. curvithorax*.

*M. gallienii*

One sample identified as *M. gallienii* showed a distinct signature of mixed ancestry in the admixture results. It was related on the mitochondrial phylogeny only to another individual also identified morphologically as *M. gallienii*, but whose nuclear data did not pass filtering. There was no published sequence available for this species, and we conclude they are true *M. gallienii*.

*M. rugulosa*

One sample identified as *M. rugulosa* showed a distinct signature of mixed ancestry in the admixture results. It was related on the mitochondrial phylogeny to references of *M. rugulosa*, so we conclude that it is a true *M. rugulosa*.

*M. vandeli*

Three samples were identified morphologically as *M. vandeli*, had mitochondrial DNA similar to a reference individual of *M. vandeli,* and had similar nuclear genome composition. They were thus considered to belong to *M. vandeli*.

*M. sulcinodis*

Our survey likely also included three samples of *M. sulcinodis*, two of which were morphologically identified as such and one as *M. scabrinodis*). The three clustered together on the MDS and formed a monophyletic COI group with the published sequence of *M. xavieri*, which is known only from Spain. The second closest published sequences on the mtDNA tree belonged to *M. sulcinodis*, so we conclude that they are most likely *M. sulcinodis*.

*M. lonae*

From the morphology-based species list, there was only one species (*M. lonae*) that we did not recover in our genetic dataset. One sample was morphologically assigned to that species. It had both mitochondrial and nuclear DNA matching *M. sabuleti*, with no signs of belonging to a different genetic group. *M. lonae* is very similar to *M. sabuleti* (Seifert 2018), and their relationships have never been studied genetically. They differ in subtle morphological traits and ecology (Seifert, 2000), which could also indicate different ecotypes or plasticity. Assessing the validity of the species status of *M. lonae* is beyond the scope of this paper and would require more extensive sampling, but in any case we conclude that this individual belongs to *M. sabuleti*.
